# Dying of thirst: Osmoregulation by a hawkmoth pollinator in response to variability in ambient humidity and nectar availability

**DOI:** 10.1101/2023.06.27.546736

**Authors:** Ajinkya Dahake, Steven Persaud, Marnesha N. Jones, Joaquin Goyret, Goggy Davidowitz, Robert A. Raguso

## Abstract

Climate-induced alterations in flowering phenology can lead to a temporal mismatch between pollinators and the availability of floral resources. Such asynchrony may be especially impactful in desert ecosystems, where flowering time and pollinator emergence are particularly sensitive to rainfall. To investigate the osmoregulation of a desert-living hawkmoth pollinator *Manduca sexta*, we sampled hemolymph osmolality of over 1000 lab-grown moths held at 20%, 50%, and 80% ambient humidity. Under starvation, the hemolymph osmolality of moths remained within a healthy range from days 1-3, regardless of ambient humidity. However, osmolality levels increase steeply from a baseline of 360-370 mmol/kg to 550 mmol/kg after 4-5 days in low and intermediate humidity and after 5 days in high humidity. Starved moths exposed to low humidity conditions died within 5 days, whereas those in high humidity conditions lived twice as long. Moths fed either synthetic *Datura wrightii* nectar, synthetic *Agave palmeri* nectar, or water, maintained osmolality within a healthy baseline range of 350-400 mmol/kg. The same was true for moths that fed on authentic floral nectars. However, moths consumed higher amounts of synthetic nectar, likely due to the non-sugar nectar constituents in the authentic nectar. Finally, simulating a 4-day mismatch between pollinator emergence and nectar availability, we found that a single nectar meal can osmotically rescue moths in dry ambient conditions. Our findings indicate that hemolymph osmolality provides a rapid and accurate biomarker for assessing both the health and relative hydration state of insect pollinators.

## Introduction

A rising concern among scientists and policymakers alike is whether pollinators can cope with fluctuations in climate and consequent changes in the availability of floral rewards (nectar and pollen) due to changes in plant blooming time (phenology) (Willmer, 2012; Straka & Starzomski, 2014; Cohen et al., 2018; Soroye et al., 2020; Stout & Dicks, 2022). These observed seasonal shifts not only alter floral phenology but also temporal aspects of pollinator biology, from emergence to migration (Forrest, 2016; Luder et al., 2018; Satterfield et al., 2020). While some studies report that both plant and pollinator phenologies have advanced by 6-10 days over the last few decades, (Bartomeus et al., 2011; Willmer, 2012; Forrest, 2015; Duchenne et al., 2020) others have documented a mismatch in flower-pollinator phenology due to changes in snowmelt time (Kudo & Ida, 2013; Kudo & Cooper, 2019). Therefore, climate change is predicted to impact plant-pollinator synchrony on a global scale, leading to re-wired plant-pollinator networks, with implications for management practices, ecosystem services, conservation, and habitat restoration (Burkle & Alarcon, 2011). Ongoing studies address such impacts on large, long-lived vertebrate pollinators such as migratory hummingbirds and bats (Fleming et al., 1993; Morales-Garza et al., 2007; Ayala-Berdon et al., 2011; Ayala-Berdon & Schondube, 2011). Here we focus on shorter-lived insects, for whom lifetime reproductive success is generally limited to a few weeks, with the potential for increased vulnerability to climate instability.

Of the several aspects of climate change that can influence plant-pollinator phenology, shifting precipitation is considered a crucial driver (Cohen et al., 2018; Henry et al., 2022). Regions that are currently dry are projected to experience more severe droughts (Trenberth, 2011) and wildfires, resulting in pollinator declines (Banza et al., 2019). Pollinators in arid habitats are especially affected by changes in precipitation because their arrival or emergence is often linked to precipitation events (Danforth, 1999; Contreras et al., 2013). The greatest natural stressors on small insects in arid climates are high temperatures and the lack of moisture or low relative humidity (RH) (Henschel & Seely, 2008). Due to their high surface area to volume ratio (Kühsel et al., 2017), insects are susceptible to water loss and must find appropriate microenvironments and sufficient food resources to guard against desiccation (Willmer, 1982; Chown et al., 2011). Unlike small insects that can inhabit humid flowers (Harrap et al., 2020; Dahake et al., 2022), cycad cones (Salzman et al., 2023), or the boundary layers of leaves (Ramsay et al., 1938; Potter et al., 2009; Woods, 2010; Potter et al., 2011; Potter et al., 2012), large insects must find alternative ways to avoid desiccation. Earlier studies show that smaller xerophilic bees (*Chalicodoma sicula*; 143 – 233 mg, *Xylocopa sulcatipes*; 376 - 422 mg) experience a water deficit during the hot and dry summer months (Willmer, 1986). In contrast, larger bees (*Xylocopa pubescens*; 536-632 mg, *Xylocopa capitata*; 1.2-1.5 g) experience water excess exacerbated by their high metabolic activity (Nicolson & Louw, 1982; Willmer, 1988; Nicolson, 1990). Therefore, the strategies used by small and large pollinators to prevent desiccation, avoid osmotic stress, and modify foraging behavior in unpredictable conditions is a topic of interest, especially with respect to monitoring pollinator health.

Recently, several papers have reviewed the theme of “pollinator health” with domestic and wild bees as focal taxa (Lopez-Uribe et al., 2020; Parreno et al., 2022). Non-destructive observation-based markers of pollinator health include monitoring body size, body weight, and foraging activity using radio frequency identification (RFID) tags, lifespan, and fecundity (Perry et al., 2015; Lopez-Uribe et al., 2020). In contrast, destructive sampling methods include sacrificing pollinators to monitor their lipid content, immunological response (immune gene expression levels), oxidative stress, Cytochrome P450 gene expression (pesticide exposure), and qPCRs for pathogen abundance, among other markers (Alaux et al., 2010; Traver & Fell, 2011; Evans et al., 2013; López-Uribe et al., 2016; Simone-Finstrom et al., 2016; Smart et al., 2016; Alaux et al., 2017; Schurr et al., 2017; Ricigliano et al., 2019; Lopez-Uribe et al., 2020). While these sampling techniques work well for social insects with large populations and a central place foraging strategy, it has been a challenge to monitor similar indices of health for insects that have solitary lifestyles, forage, or disperse over long distances. Although much attention has been given to bees, the health impacts on other insect pollinators, including moths (Lepidoptera) are less commonly addressed (Stevenson et al., 2022).

Moths outnumber bees by an order of magnitude in species richness but likely go unnoticed due to their public connotation as agricultural or stored product pests, as well as their predominantly nocturnal niche. Despite this, moths serve as valuable pollinators and outcrossing agents (Devoto et al., 2011), especially to guilds of night-blooming flowers (MacGregor et al., 2015; Hahn & Brühl, 2016), particularly in urban environments (Ellis et al., 2023). Among the best-studied group of moths as pollinators are the sphinx moths or hawkmoths (Sphingidae), most of which are large-bodied animals (1-3 g) that can be highly effective pollinators (Oliveira et al., 2004; Johnson et al., 2017), especially over distances not traversed by most bees (Skogen et al., 2019). The Carolina Sphinx (tobacco hornworm) moth, *Manduca sexta* is a large nocturnal hawkmoth (2 g) found in many different habitats across the Americas. The ecology of *Manduca sexta* has been extensively studied in the Sonoran Desert of Arizona, USA (Raguso & Willis, 2005; Bronstein et al., 2007; Alarcón et al., 2008; Riffell et al., 2008; Bronstein et al., 2009; Wilson et al., 2018; Wilson et al., 2019; Johnson et al., 2021). The Sonoran Desert spreads over 300,000 sq. km and spans 2 countries and 5 states (Comus et al., 2015). In arid southwestern USA, annual precipitation is projected to decrease, with concomitant projected increases in droughts and wildfires (Easterling, 2017; Overpeck & Udall, 2020; *NOAA Drought Task Force Report on the 2020–2021 Southwestern U.S. Drought*, 2021). Recent efforts have elucidated the major role that ambient humidity plays in the life histories of these nocturnal pollinators. Light trap data collected from the Santa Rita Experimental Range (SRER) near Tucson, AZ show that hawkmoth occurrence peaks with the onset of monsoon rains (July and August) in the Sonoran Desert (Contreras et al., 2013). In this otherwise dry habitat, the arrival of monsoon rains can increase relative humidity (RH) from 20 to 80%. Thus, increased ambient humidity is tracked by hawkmoth abundance, through synchronized adult emergence or immigration (Contreras et al., 2013), as shown previously for desert bees (Danforth, 1999).

Lab experiments show that moths survive twice as long in a background RH of 60-80% than in a background of 20-40% RH. Thus, there should be a strong selection for moths to perceive differences in ambient RH. Indeed, the antennae of *M. sexta* are equipped with specialized hygrosensory sensilla, the underlying neurons of which respond to tiny fluctuations in ambient humidity. Minute rates of change (0.23%RH/sec and 0.30%RH/sec) are sufficient to trigger a spike in the antennal moist- and dry-sensing neurons, respectively (Dahake et al., 2022). Accordingly, moths should be equipped to show clear behavioral responses to differences in ambient RH. In wind-tunnel choice assays, *M. sexta* moths significantly favor the more humid air stream of the tunnel, even if the difference between the two sides is less than 8% RH (Wolfin et al., 2018). Moreover, adult moths regulate their consumption of nectar in different ambient humidity levels by showing a preference for dilute nectar (12% sucrose) or water (0% sucrose) in low RH and concentrated nectar (24% sucrose) in high RH (Contreras et al., 2022). However, despite differences in preference, moths appear to use compensatory feeding to maintain a constant overall energy intake (Contreras et al., 2013; Contreras et al., 2022), emphasizing the physiological balance between sometimes competing caloric and osmoregulatory demands. Clearly, ambient conditions can modify pollinator behavior and affect their survivorship.

Field sampling studies provide snapshots of pollinator health during specific environmental conditions or periods. What is needed are longitudinal studies that track pollinator physiology under variable environmental conditions, establishing baseline performance data that can lead to a more predictive framework for tracking pollinator health (Brochu et al., 2020). How do we go beyond monitoring seasonal changes in community structure and population dynamics, to quickly monitor the physiological state of pollinators? What is the degree of physiological plasticity shown by pollinators and how does their internal physiological state reflect the choice of flowers they visit? We address some of these questions using *Manduca sexta* as a model organism. Earlier work on large bees suggested that larger pollinators, such as hawkmoths, might experience water excess due to their intense metabolic activity (Heinrich & Raven, 1972; Heinrich, 2013). Instead, we found that *Manduca sexta*, despite being one of the largest insect pollinators, experiences severe water loss in dry conditions. We employed a laboratory osmometer to establish baseline hemolymph concentrations of starved and well-fed moths held in dry (20% RH, vapor pressure deficit (VPD) = 2.4 kPa), intermediate (50% RH, VPD = 1.5 kPa), and humid environments (80% RH, VPD= 0.6 kPa). Although hemolymph osmolalities of some desert insects have been documented previously, there is a lack of systematic baselines as a reference to compare individuals of different ages and physiological states (Nicolson, 1980, 1990). We evaluate the osmoregulatory capacity of *M. sexta* hand-fed with two nectar diets (*Agave palmeri, Datura wrightii*) and water, which are available in their natural habitat in the Sonoran Desert. We compare the differences in feeding amounts of authentic vs. synthetic nectar and their impact on moth hemolymph osmolality. Finally, we test the impact of an experimentally simulated 4-day mismatch between pollinator emergence and nectar availability on moths’ ability to rescue themselves from osmotic stress. Our findings suggest that hemolymph osmolality could serve as a rapid and accurate biomarker for the health (dehydrated vs. satiated state) of pollinators in their natural habitats.

## Materials and methods

### Animals

Continuously breeding laboratory colonies of *M. sexta* (Contreras et al., 2022) were maintained in parallel in the Raguso lab at Cornell University and the Davidowitz lab at the University of Arizona, Tucson. Larvae were reared to maturity on an artificial diet, under conditions described by Davidowitz et al. (2003). Pupae were separated by sex and were transferred to incubators with different experimental treatments for relative humidity (RH) so that adult moths could eclose directly into the assigned RH treatment.

### Experimental setup for the three humidity levels

Three environmental incubators (Thermo Scientific, Inc.) were maintained at 24°C and programmed to a 16:8 light: dark cycle. The incubators were set to operate at 20%, 50%, or 80% RH (with corresponding vapor pressure deficits of air in kPa [VPD] of 2.4, 1.5, and 0.6, respectively). At Cornell, the following steps were taken to produce stable differences in incubator RH. To maintain 80% RH, a 2L humidifier (Vicks, Inc.) was placed inside the incubator, with an electronic relay controlled using a Raspberry Pi device that received instantaneous RH values from an Adafruit SHT31 temperature and humidity sensor also placed inside the incubator. This created a closed-loop system to maintain a stable RH of 80±2% (mean ± SD). For 50% RH, two water buckets with approximately 2L water each were placed inside the incubator, and for 20% RH, an automatic dehumidifier from YouFu, Inc. set at 20% RH level was used. At Arizona (with drier ambient RH conditions in the laboratory), to achieve 80% RH, we used small humidifiers connected to the incubators through hoses, which remained engaged without any feedback device. The humidifier was refilled twice a day (morning and night). For 50% RH, we placed a small tray of water, enough to fully cover the bottom, in the incubator (this was usually done every morning and night). There were no alterations done to maintain 20% RH because the ambient RH typically ranged between 20 - 25%. For the rescue experiment a Percival (I-36VL) automatic incubator was used and set at 24℃ and 80% RH. All other humidities were maintained using the methods described above.

### Hemolymph osmolality

Appropriately aged moths were removed from the incubator and anesthetized in a −20°C freezer for 5-10 mins. Anesthetized moths were weighed (fresh mass to 0.001g) and taken to the fume hood for descaling the “neck” of the moth (the intersegmental membrane between head and thorax), using pressurized air and/or a kitchen scouring pad (3M, Inc.). The dorsal surface of the neck was punctured roughly at the midline with an insect pin. The moth’s neck was stretched ventrally, and pressure was applied to the thorax with the thumb to collect 10µl of oozing hemolymph in a 10ul microcapillary glass tube (Fisherbrand, Inc.). In cases where the neck did not yield enough hemolymph, pressurized air was blown through the hole in the neck such that hemolymph accumulated in the last abdominal segment, which was then punctured as described, using the insect pin. Hemolymph solute concentration for the same animal did not differ between the neck and abdomen (results not shown). The hemolymph sample was pipetted onto a small filter paper disc, then placed within the sample holder of a Wescor 5600 Vapor Pressure Osmometer for analysis. Osmolality is a measurement of the number of solute particles in a kg of solvent (In this case moth hemolymph) and is independent of temperature. Osmolality can be expressed as mOsmol/kg of solvent or the SI unit mmol solute/kg solvent. A higher osmolality indicates more solutes, and in the context of this study, a higher osmolality indicates greater desiccation of the moth. Seawater is on average 1000 mmol/kg, human blood ranges between 275-295 mmol/kg and deionized water is 0 mmol/kg. The osmometer was calibrated either once each week or after every 5-10 readings using Opti-mole osmolarity standard solutions of 100 mmol/kg, 290 mmol/kg, and 1000 mmol/kg provided by Wescor. Osmolality data were collected for as long as 10 µl hemolymph could be extracted from moths under the different ambient humidity environments. Moths were only used once for osmolality measurements and the same methods were followed for all experiments described in this manuscript. After hemolymph extraction, moths were stored in glassine envelopes and were euthanized in a −20°C freezer.

### Water loss experiment

Insects are particularly prone to desiccation risk (Chown et al., 2011). The fresh mass of adult moths was used as a proxy to determine active vs passive water loss from adult moths. Pairs of male and female *Manduca sexta* pupae were isolated from the lab colony and placed in separate nylon mesh cages (35 x 35 x 60 cm; Bioquip Products, Inc.) inside climate-controlled incubators. On the day when moths eclosed from the pupae, one individual of each sex was immediately freeze-killed and returned to the cage. The dead moth served as a control to account for passive evaporation through the cuticle occurring in the living moth. Both the dead and the living moths were weighed daily to 0.01g using a digital balance (Mettler Toledo) from the day of eclosion (day 0) for as long as moths survived at each ambient RH level. We used 10 moths (for each living and dead) for the three ambient RH levels. Accounting for both sexes, a total of 120 moths were used in this experiment.

### Starvation experiment

To establish baseline values for hemolymph osmolality under different ambient RH levels, male and female pupae were placed in separate nylon mesh cages in climate-controlled incubators. All emerging moths were starved (unfed) in this experiment. Moths that still had wet wings (freshly eclosed) were considered 1 day old the following day. At a specific age (days post-eclosion), moths were removed from the incubator to draw hemolymph samples. New pupae were introduced into the cage until a sample size of 10 moths per day was achieved. Although half of the total number of moths survived beyond the day at which 50% mortality was reached, these moths rarely yielded enough hemolymph for a reliable osmolality reading. Reliable osmolality data were obtained up to days 5, 6, and 8 for moths held at 20%, 50%, and 80% RH, respectively.

### The liquid diet used in the feeding experiments

The flight activity of *Manduca sexta* in the Sonoran Desert overlaps with the monsoon season and the flowering phenology of *Agave* and *Datura* plants, which are their primary nectar sources (Fig. 1A) (Raguso et al., 2003; Alarcón et al., 2008; Bronstein et al., 2009; Contreras et al., 2013). To investigate the ability of male *M. sexta* moths to maintain hemolymph osmolality under different diets and relative humidity (RH) conditions, two synthetic nectar mixtures were created. The first synthetic mixture replicated the nectar sugar composition of *Datura wrightii* flowers, consisting of 337.5 mg/ml of glucose, 276.79 mg/ml of fructose, and 1589.29 mg/ml of sucrose (Riffell et al., 2008). This mixture was the first of three diets and had a sugar concentration of approximately 22% with a high proportion of sucrose (Fig. 1B). The second synthetic mixture replicated the nectar sugar composition of *Agave palmeri* flowers, consisting of 651.29 mg/ml of glucose, 547.56 mg/ml of fructose, and 1.95 mg/ml of sucrose (Riffell et al., 2008). This mixture was the second of three diets and had a sugar concentration of approximately 12% with a high proportion of hexose sugars (Fig. 1C). The last diet consisted of deionized water and therefore had 0% sugar (Fig. 1D).

### Ad Libitum Feeding of Moths

Each climate-controlled incubator consisted of three nylon mesh cages, where moths eclosed separately and were fed one of three diets ad libitum. Feeding was conducted on laboratory benchtops outside of the incubators. Three separate feeding experiments were conducted: one with synthetic nectar, one with authentic nectar, and a rescue experiment in which moths were fed synthetic nectar or water. Males and females did not differ in their osmolality curves; therefore, the feeding experiments were only conducted on males. The same feeding protocol was followed for all subsequent experiments. Droplets of the appropriate liquid diet (50µl) were pipetted onto Parafilm wells. Moths were removed from the incubators and held by hand along the strongly reinforced costal vein of the forewing. The moth’s proboscis was extended using an insect pin and placed into a liquid droplet. Once a droplet was fully consumed, the proboscis was moved to a second droplet, and this process was repeated until the moth was fully satiated. Any remaining liquid was pipetted up and measured (to 1 µl). Each moth’s proboscis was extended and placed into the droplet at least three times to ensure it was fully satiated. Fed moths were returned to their respective incubators and maintained at different RH levels. Moths were fed every day except for the day of hemolymph extraction, to prevent recently imbibed liquid from confounding osmolality values taken on that day.

**Fig. 1.**
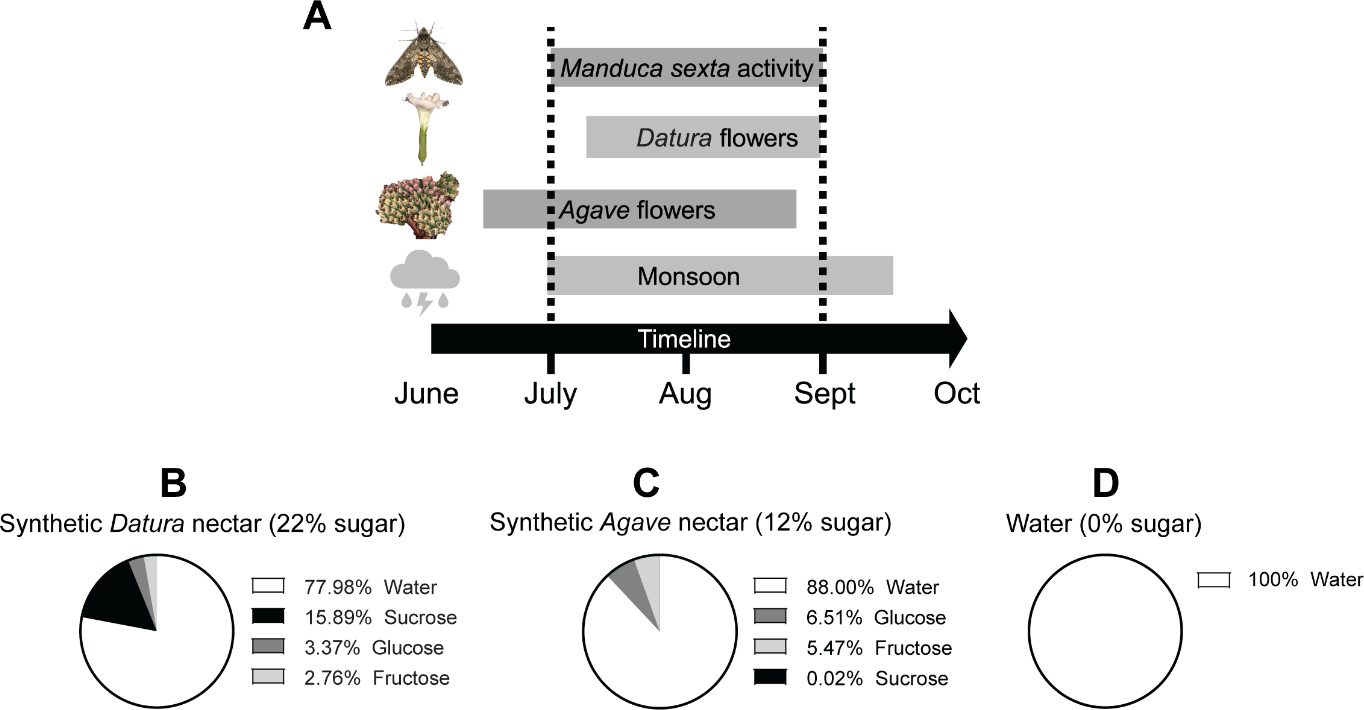
An outline justifying the use of the three diets and the ambient humidity used in our experiments throughout this study. (A) shows the overlapping timeline between Monsoon, *Manduca sexta* activity, and the flowering period of two plants that serve as major sources of nectar meals in the Sonoran Desert during the months of July-Sept. The ambient humidity during Monsoon season in Tucson, Arizona, USA ranges between 20% to 80% RH (See fig. 1 in Contreras et al. (2013)). (B-D) Pie charts show the nectar composition of the three liquid diets as total dissolved solids: synthetic *Datura wrightii* nectar, synthetic *Agave palmeri* nectar, and deionized water (After Riffell et al. (2008)). The proportions of the different sugar combinations are color coded.

For the *synthetic nectar feeding* experiment, male *M. sexta* pupae were placed in the incubators maintained at 20%, 50%, or 80% RH. Moths were fed ad libitum as described above from day 1-7 post-eclosion, and their osmolality was measured each day until day 8. However, these are not repeated measures because once osmolality was recorded, the moth was sacrificed, and a new moth was introduced to the treatment-appropriate cage. We aimed for a sample size of 5-8 moths for each day post-eclosion. Thus, the data shown in Fig. 4 required us to destructively sample 374 moths across the three diets and ambient RH levels. Although nectar-fed moths typically survive beyond day 8, we terminated the experiment here to compare osmotic concentrations with the starved moths that had survived a maximum of 8 days.

For the *authentic nectar feeding* experiment, moths were fed either *Datura wrightii* flower nectar (with a sugar concentration of approximately 24%) or *Agave palmeri* nectar (sugar conc. ∼12%). This nectar was previously collected from newly opened flowers and was frozen for storage at −20°C. *Datura wrightii* nectar was collected from greenhouse-grown plants at Cornell University and *Agave palmeri* nectar was collected from natural populations growing in the SRER (see above) in Arizona. In this experiment, moths were fed ad libitum for 4 days, and their hemolymph was extracted on day 5. Day 5 was selected as a single time point for measurement because in the starvation experiment, hemolymph osmolality of moths reached lethal levels at this point (see Fig. 2C, D). Osmolality values of 5 to 9 moths were obtained for each combination of RH and diet, with each moth being 5 days old.

**Fig. 2.**
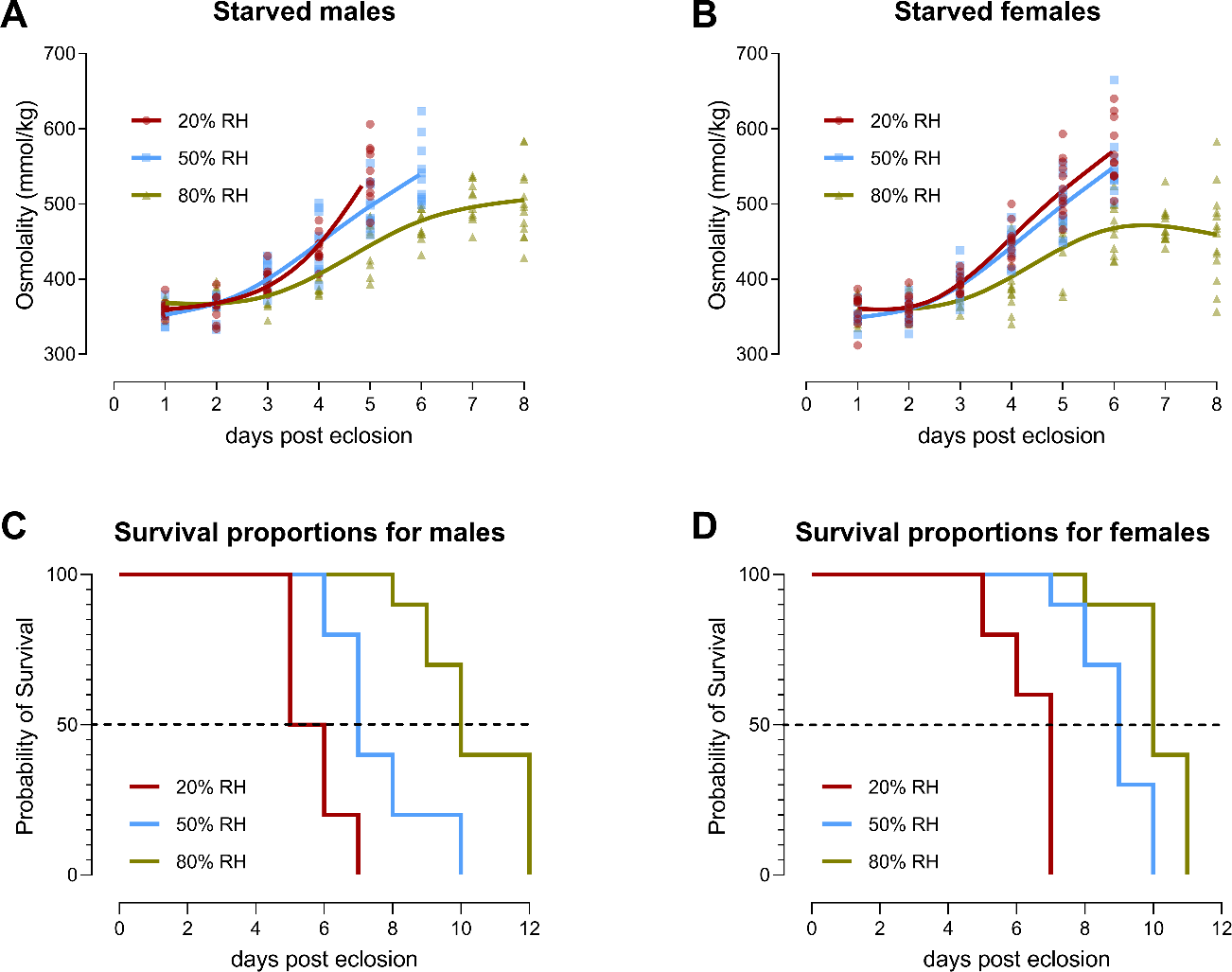
The effect of starvation on the hemolymph osmolality and longevity of *Manduca sexta* held at 20%, 50%, and 80% ambient RH. (A&B) Starvation-induced impact on blood osmolality (mmol/kg) of n=10 males and females each held at three different levels of ambient RH. Data points are fitted with a 4-knot smoothing cubic spline curve. Note that male moths yield hemolymph by day 5 in 20% RH and day 6 in 50% RH, while females in 20% and 50% RH yield hemolymph only until day 6 post eclosion. (C &D) Kaplan-Meier survival curves of male and female moths held at the three ambient RH levels. The dashed line indicates a 50% survival rate. Lines are color-coded to represent the ambient humidity level.

For the *rescue* experiment, we evaluated whether a single meal could restore water balance in moths. We starved newly eclosed moths for three days, fed the moths *ad libitum* on day four, and measured hemolymph osmolality on day five. As a control, we had a separate group of moths that were left unfed, and their hemolymph samples were taken on either day 3 or day 5. The experiment with the authentic *Agave* nectar and the rescue experiment were conducted at the University of Arizona, in the Davidowitz lab. All other experiments were conducted at Cornell University, in the Raguso lab.

### Statistics

All statistics were performed in either GraphPad Prism (version 9.5.1) or R Studio (version 4.1.1) The osmolality curves of the starved moths in Fig. 2 were compared using a two-way ANOVA with fixed effects for day, ambient RH, and their interaction. Model effects were tested with an F-test, followed by post hoc multiple comparisons using Tukey’s method using the “emmeans” package in R. The Kaplan-Meier survival curves for moths in the different ambient RH levels were compared using the log-rank Mantel-Cox test. Linear regressions were used to analyze the data from the *water loss* experiment and the *synthetic nectar-feeding* experiment, with an F-test conducted across the different humidity levels. For a comparison between the osmolality of starved vs fed moths, we hypothesized that the fed moths would show lower osmolality (less dehydrated) compared with the starved moths based on the results of previous experiments. We performed individual comparisons using an unpaired t-test with Welch correction. The *authentic nectar*-*feeding* experiment and the feeding amounts in the *rescue* experiment were analyzed using either a one-way ANOVA or a Kruskal-Wallis test depending on data distribution and variance structure. These tests were followed by Fisher’s LSD test for individual comparisons. The rescue experiment feeding amounts were tested using an ANOVA, followed by a t-test for individual comparisons. The osmolality data were compared with a one-tailed one-sample t-test where the mean osmolality for each treatment was tested against the mean baseline osmolality.

## Results

### The osmolality of starved moths

We predicted that hemolymph osmolality should increase as the moths become more dehydrated. Our results are consistent with this prediction. The hemolymph osmolality of male and female moths increases with age when they are starved, but the rate of increase depends on the ambient humidity. Initially, the average hemolymph osmolality of moths on the first and second day after eclosion, regardless of humidity level, is around 350-360 mmol/kg (Fig. 2A & B). However, starting from the third day of starvation, the osmolality of moths in 20% and 50% relative humidity (RH) increases rapidly and deviates significantly from the moths in 80% RH (Table 1). This difference in osmolality is maintained on the fifth and sixth day of starvation for both males and females. Moths in 80% RH experience a gradual increase in osmotic concentration. By the fifth day of starvation, the hemolymph osmolality of moths held at 20% RH exceeds 500 mmol/kg, followed by moths held at 50% RH at around 500 mmol/kg, and moths in 80% RH at around 450 mmol/kg. The moths in 20% RH and 50% RH die by the sixth day after eclosion when the hemolymph concentration peaks over 550 mmol/kg. However, the moths held at 80% RH never exceed 550 mmol/kg, even by the eighth day, but fail to yield enough hemolymph for further analysis beyond day 8 (Fig. 2A & B).

**Table 1.** Multiple comparisons of the two-way ANOVA followed by Tukey’s post hoc test for osmolality of starved moths in the three ambient humidity levels. *P*-values for days 1 and 2 are not shown because none of the combinations (ANOVA) were statistically significant.

|  |  |  |  |  |  |
| --- | --- | --- | --- | --- | --- |
| Day | Multiple comparisons between different levels of RH | Sex | d.f. | *t* ratio | *P*-value |
| 3 | 20% vs 50% | M | 135 | -0.840 | 0.6789 |
|  |  | F | 168 | -0.022 | 0.9997 |
|  | 50% vs 80% | M | 135 | 2.785 | **0.0168** |
|  |  | F | 168 | 1.555 | 0.2682 |
|  | 20% vs 80% | M | 135 | 1.945 | 0.1303 |
|  |  | F | 168 | 1.533 | 0.2780 |
| 4 | 20% vs 50% | M | 135 | -0.311 | 0.9481 |
|  |  | F | 168 | 0.451 | 0.8940 |
|  | 50% vs 80% | M | 135 | 4.324 | **0.0001** |
|  |  | F | 168 | 4.734 | **<0.0001** |
|  | 20% vs 80% | M | 135 | 4.013 | **0.0003** |
|  |  | F | 168 | 5.205 | **<0.0001** |
| 5 | 20% vs 50% | M | 135 | 3.548 | **0.0015** |
|  |  | F | 168 | 2.150 | 0.0831 |
|  | 50% vs 80% | M | 135 | 4.431 | **0.0001** |
|  |  | F | 168 | 3.854 | **0.0005** |
|  | 20% vs 80% | M | 135 | 7.979 | **<0.0001** |
|  |  | F | 168 | 6.029 | **<0.0001** |
| 6 | 20% vs 50% | F | 168 | 0.766 | 0.7243 |
|  | 50% vs 80% | F | 168 | 6.038 | **<0.0001** |
|  | 20% vs 80% | F | 168 | 6.822 | **<0.0001** |

*Manduca sexta* lifespan (males and females) was significantly impacted by the ambient humidity at which they were held (Fig. 2C & D; Log-rank Mantel-Cox test for males: *Χ^2^*=28.80, d.f.=2, *P*<0.0001; females: *Χ^2^*=33.99, d.f.=2, *P*<0.0001). In general, moths died at a younger age when kept at lower levels of humidity when compared with higher levels (Table 2), and there is strong evidence of a difference in lifespan among the three levels of ambient relative humidity (RH) (Table 2). The order of moth lifespan, for both males and females, was as follows: 20% RH < 50% RH < 80% RH.

**Table 2.** Comparison of lifespan of starved male and female *M. sexta* held at different ambient RH levels.

|  |  |  |  |  |  |  |  |
| --- | --- | --- | --- | --- | --- | --- | --- |
| Ambient RH | Sex | Lifespan in days (mean ± s.d.) | 50% mortality (days) | Lifespan comparisons between different levels of RH | d.f. | Mantel-Cox test $\chi^2$ | *P*-value |
| 20% | M | 5.7 ± 0.8 | 5 | 20% vs 50% | 1 | 9.72 | **0.0018** |
| | F | $6.4 \pm 0.8$ | 6 | | 1 | 15.65 | **0.0001** |
| 50% | M | $7.6 \pm 1.3$ | 7 | 50% vs 80% | 1 | 9.54 | **0.0020** |
| | F | $8.9 \pm 0.9$ | 10 | | 1 | 8.34 | **0.0039** |
| 80% | M | $10.4 \pm 1.4$ | 10 | 20% vs 80% | 1 | 21.00 | **0.0001** |
| | F | $10.2 \pm 0.9$ | 11 | | 1 | 18.31 | **0.0001** |

### Active vs passive weight loss in adult moths

We investigated the impact of ambient relative humidity (RH) levels (20%, 50%, and 80% RH) on weight loss in male and female *Manduca sexta*. To determine whether weight loss is primarily due to passive evaporation or active metabolic processes, we compared the weight loss of living moths with that of equally aged dead moths. The results showed that living moths lost more weight compared to their paired dead moths undergoing passive evaporation, regardless of the ambient humidity level (Fig. 3). However, a significant difference in weight loss was observed for both male and female moths held at higher ambient humidity (Table 3). Moths in dry conditions did not show significant differences in weight loss compared to the passively evaporating dead moths. Moths held at 20%, 50%, and 80% RH lost weight in the order of 20% RH > 50% RH > 80% RH. The slopes of the regression lines for weight loss were not significantly different among males at different RH levels (*F*=0.9131, d.f.=2 and 191, *P*=0.4030), but were among females (*F*=4.111, d.f.=2 and 98, *P*=0.0178).

**Fig. 3.**
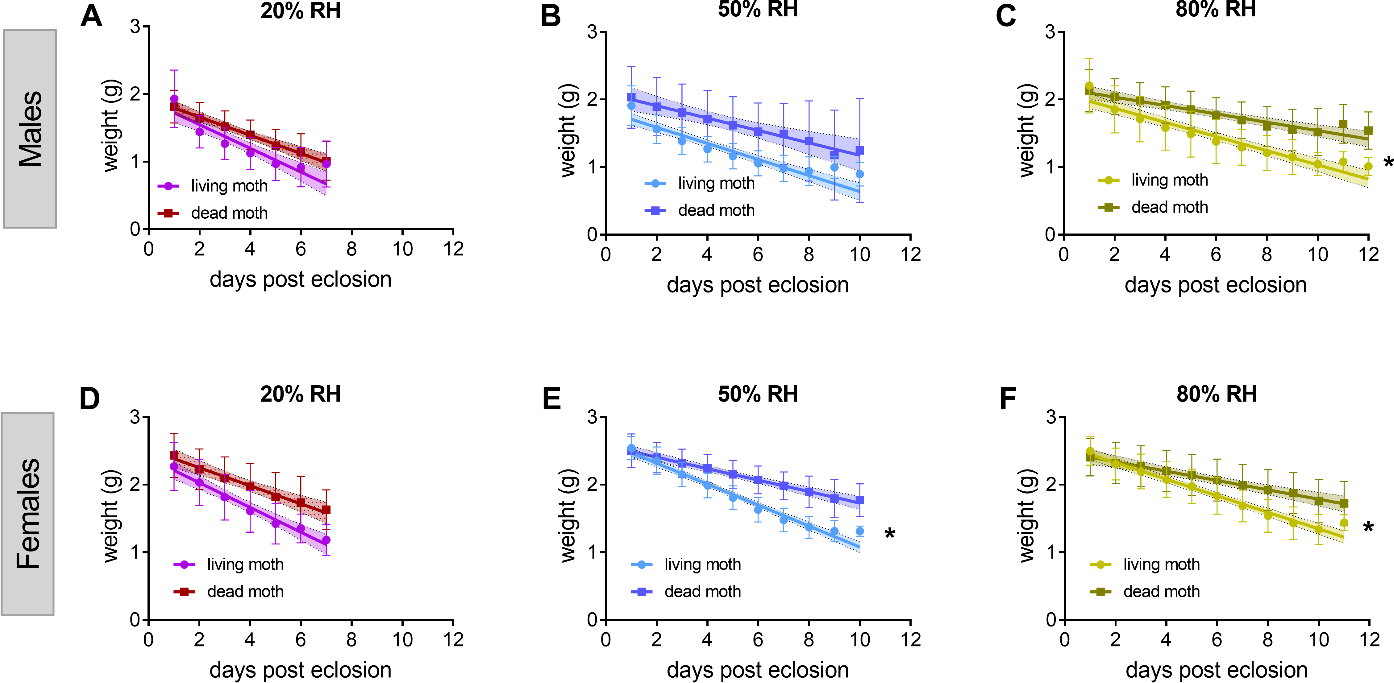
Weight loss in fresh mass to 0.01g of starved adult moths in 20%, 50%, and 80% ambient RH. The weight loss of active vs. passive males (A-C) and females (D-E) *M. sexta* weight loss is compared with passively evaporating paired dead moths in the three ambient RH levels. Data are color-coded by the ambient RH and the condition of the moth (living vs. dead). Points show the mean and error bars show the SD for each day and are fitted with linear regression lines. Data are repeated measures from n=10 moths for both conditions (living and dead) for each ambient RH. * Indicates a significant difference in the slope of the two linear fits.

**Table 3.**
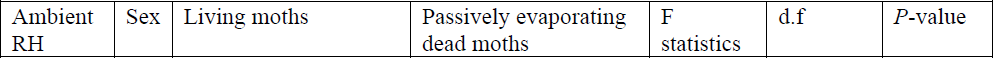
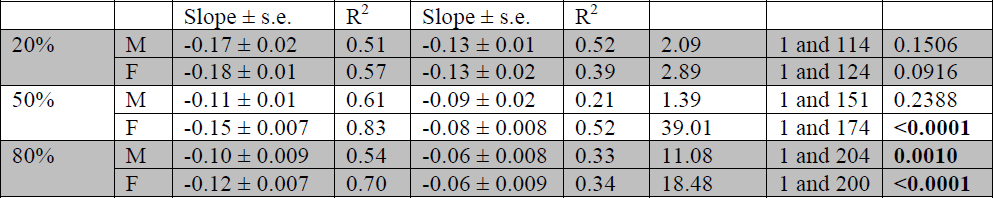
Summary statistics of the linear regression fit for weight loss in the living and the dead *M. sexta* in the three ambient RH levels.

### The osmolality of moths fed on synthetic nectar or water

Since the trends in osmolality were similar between males and females (Fig. 2), we conducted this experiment exclusively on males. As moths age, their consumption of synthetic nectar or water increases 2- to 4-fold from days 1 to 7 after eclosion. This consumption pattern remained consistent across the three humidity levels, with the following order: water < synthetic *Datura* nectar < synthetic *Agave* nectar (Fig. 4A-C). However, the amount of synthetic nectar consumed did not differ among moths in different ambient RH levels (Fig. 4A-C). Nevertheless, moths in 20% RH consumed significantly higher amounts of water than did moths in 50% RH (*F*=5.318, d.f.=1 and 305, *P*=0.021) and in 80% RH (*F*=12.22, d.f=1 and 311, *P*<0.001). The hemolymph osmolality of fed moths remained within the range of 350-400 mmol/kg, which was in stark contrast to the osmolality ranges observed in the starvation experiment (Fig. 2). Furthermore, there was no evidence of a difference in the osmolality ranges for moths that fed on different diets (Fig. 4D-F). However, there were several occurrences of the moths in water-fed group that either did not yield enough hemolymph or died before day 8. This was prominent in 20% ambient RH (Fig. S1). None of the nectar-fed moths died before day 8. The dead moths were replaced by introducing new pupae to achieve sufficient sample sizes for each treatment group.

**Fig. 4.**
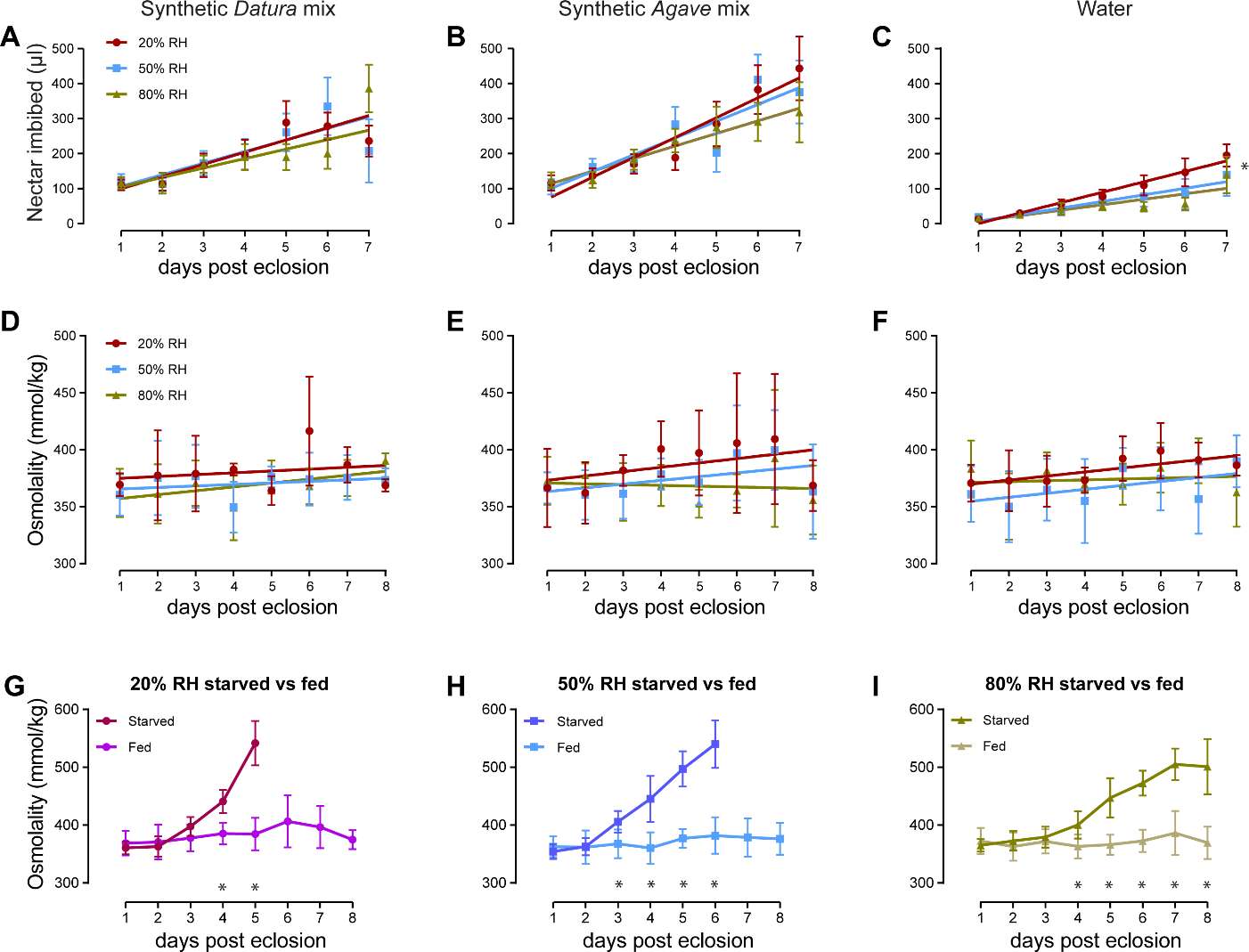
A comparison between the dehydrated and synthetic nectar-fed moths. (A-C) the amount of synthetic nectar or water imbibed (mean ± SE) by moths in the three different ambient RH levels. (D-F) the corresponding osmolality of moths fed on either of the three diets (shown above). A single day represents a sample size of n=5-7 moths. Data points show the mean and error bars show S.D. and both are color-coded to represent the ambient humidity level. Points are fitted with linear regressions. (G-I) Comparisons of the osmolality of starved moths from Fig. 2 and combined osmolality of all nectar-fed moths (n=15-17) from panels (D-F). * Indicates a significant difference for a given day (P<0.05). These experiments were performed only on the males.

Comparisons between the osmolality of starved (Fig. 2) vs. nectar-fed moths (combined across diets) showed clear effects of animal internal state on hemolymph osmolality (Fig. 4G-I). In general, for days 1 to 3 post eclosion, moths did not differ in their hemolymph osmotic concentration between the starved vs. satiated state (Table 4). Osmolality difference on day 4 and beyond between the two internal states was drastic and consistent across the three ambient RH levels (Table 4).

**Table 4.** Summary statistics of hemolymph osmolality comparisons between starved and nectar-fed moths across days post eclosion. Comparisons are conducted using unpaired t tests with Welch correction.

|  |  |  |  |  |  |  |  |  |
| --- | --- | --- | --- | --- | --- | --- | --- | --- |
| Ambient RH | Days post eclosion | Starved moth osmolality (mmol/kg) |  | Nectar-fed moth osmolality (mmol/kg) |  | *t* ratio | df | *P*-value |
| | | Mean ± s.d | N | Mean ± s.d | N | | | |
| 20% RH | 1 | 360.8 ± 11.27 | 10 | 368.7 ± 21.05 | 15 | 1.220 | 22.23 | 0.2351 |
| | 2 | 363.0 ± 17.70 | 10 | 370.7 ± 30.09 | 15 | 0.807 | 22.77 | 0.4277 |
| | 3 | 397.5 ± 16.60 | 10 | 377.7 ± 23.00 | 15 | 2.494 | 22.79 | **0.0203** |
| | 4 | 440.9 ± 19.82 | 10 | 385.4 ± 18.74 | 15 | 7.000 | 18.66 | **<0.0001** |
| | 5 | 541.8 ± 38.35 | 10 | 384.4 ± 28.04 | 15 | 11.14 | 15.31 | **<0.0001** |
| 50% RH | 1 | 354.3 ± 12.79 | 10 | 362.6 ± 18.09 | 15 | 1.343 | 22.86 | 0.1924 |
| | 2 | 362.8 ± 14.86 | 10 | 361.8 ± 28.85 | 15 | 0.113 | 21.95 | 0.9106 |
| | 3 | 405.7 ± 18.49 | 10 | 367.5 ± 24.73 | 15 | 4.408 | 22.60 | **0.0002** |
| | 4 | 445.4 ± 39.74 | 10 | 360.3 ± 27.32 | 16 | 5.944 | 14.35 | **<0.0001** |
| | 5 | 497.1 ± 30.65 | 10 | 377.0 ± 16.06 | 15 | 11.39 | 12.33 | **<0.0001** |
| | 6 | 540.1 ± 41.11 | 11 | 381.8 ± 31.51 | 15 | 10.68 | 18.08 | **<0.0001** |
| 80% RH | 1 | 365.1 ± 10.85 | 10 | 372.6 ± 22.65 | 15 | 1.106 | 21.36 | 0.2811 |
| | 2 | 372.7 ± 16.74 | 10 | 363.0 ± 24.49 | 15 | 1.176 | 22.96 | 0.2516 |
| | 3 | 379.4 ± 18.03 | 10 | 371.7 ± 20.86 | 15 | 0.977 | 21.31 | 0.3394 |
| | 4 | 400.5 ± 23.64 | 10 | 363.1 ± 20.62 | 17 | 4.155 | 16.95 | **0.0006** |
| | 5 | 447.2 ± 34.12 | 10 | 366.1 ± 17.37 | 15 | 6.937 | 12.15 | **<0.0001** |
| | 6 | 472.5 ± 21.63 | 10 | 372.7 ± 19.22 | 16 | 11.93 | 17.51 | **<0.0001** |
| | 7 | 505.1 ± 27.15 | 11 | 386.3 ± 37.90 | 16 | 9.493 | 24.92 | **<0.0001** |
| | 8 | 501.0 ± 47.69 | 13 | 369.4 ± 27.84 | 15 | 8.741 | 18.73 | **<0.0001** |

### The osmolality of authentic nectar-fed moths

A different set of moths were fed either authentic *Datura wrightii* or *Agave palmeri* nectar from days 1 through 4, and their osmolality were measured on day 5. Consistent with the previous experiment using synthetic *Datura* nectar, moth consumption increased by approximately two-fold with age. There was no evidence of differential consumption of *Datura* nectar among the three ambient RH levels (slope: *F*=0.1756, d.f.=2&6, *P*=0.8431). For the *Agave* nectar-feeding group, moths in 50% RH consumed the highest amount of *Agave* nectar on day 1, which gradually decreased as the moths aged. Moths in 20% RH gradually consumed more nectar as they aged, while moths in 80% RH consumed less as they aged.

There was a significant difference in the hemolymph osmolality of the *Datura* nectar-fed moths in the different ambient RH levels (One-way ANOVA, *F*=3.678, *P*=0.045). Specifically, the osmolality of *Datura* nectar-fed moths in 20% RH was higher than that of moths in 50% RH (Fisher’s LSD test: *t*=2.27, d.f=18, *P*=0.035) and 80% RH (Fisher’s LSD test: *t*=2.26, d.f=18, *P*=0.036). In the *Agave* nectar-feeding group, there was no evidence of a difference in the hemolymph osmolality of moths among different ambient RH levels (One-way ANOVA, *F*=0.2972, *P*=0.7461).

We compared the day 5 osmolality of the authentic nectar-fed moths with the day 5 osmolality of the starved and synthetic nectar-fed moths (shown in Figs. 2 & 4). In all cases, irrespective of the diet, moth hemolymph osmolality of fed moths was significantly lower than that of starved moths (Fig. 4G, H and Table A1). In conclusion, these data suggest that moths can maintain blood-water homeostasis with both high sucrose-rich (*Datura*) and high hexose-rich nectar (*Agave*).

### Rescue experiment

To evaluate whether a single nectar meal of any of the three diets can restore hemolymph osmotic concentration to baseline (as observed in Fig. 4), we starved moths for 3 days, fed them on day 4, and took blood samples on day 5. As a control, we had a separate group of moths that were starved for 2 or 4 days and osmolality measurements were taken on days 3 and 5, respectively. As opposed to the trends observed in the synthetic nectar feeding experiment (Fig. 4), here moths ingested higher amounts of synthetic *Datura* nectar than the dilute synthetic *Agave* nectar. However, this difference was not significant in either of the ambient RH levels (t-tests; 20%RH: *t*=1.70, *P*=0.11; 50%RH: *t*=0.55, *P*=0.58; 80%RH: *t*=0.07, *P*=0.93). Moths did feed high amounts of *Datura* nectar compared to water in 20% RH (t-test, *t*=2.79, *P*=0.01). No other significant differences in feeding amounts were observed between the different diets. For the osmolality of unfed moths on day 3, the trends were consistent with our previous experiment (Fig. 2). In all three humidity levels, the osmolality of day 3 unfed moths was well within the baseline range of 350-400 mmol/kg observed in figures 4 and 5 (Fig. 6, Table 5). The mean osmolality of moths in other conditions was compared against a baseline mean of 359.6 mmol/kg (an average of the day 3 unfed controls). Day 5 starved moths showed significantly higher than baseline hemolymph concentration in only 20% ambient RH (Table 5). A single meal of synthetic *Datura* nectar was sufficient to restore osmotic balance in 20% RH ambient conditions, but synthetic *Agave* nectar and water were not. Nevertheless, nectar or water fed moths showed significantly lower osmolality compared with day 5 starved moths (One-tailed t-test: synthetic Agave-fed: *t*= 1.98, *P*=0.032; synthetic Datura-fed: *t*=3.10, *P*=0.007; water-fed: *t*=2.01, *P*=0.032). Starved moths held at 50% and 80% RH were able to maintain osmolality within a baseline range (Fig. 6, Table 5), implying that they did not need to be rescued. In summary, a single ad-libitum nectar meal can mitigate osmotic stress in desiccating environments.

**Fig. 5.**
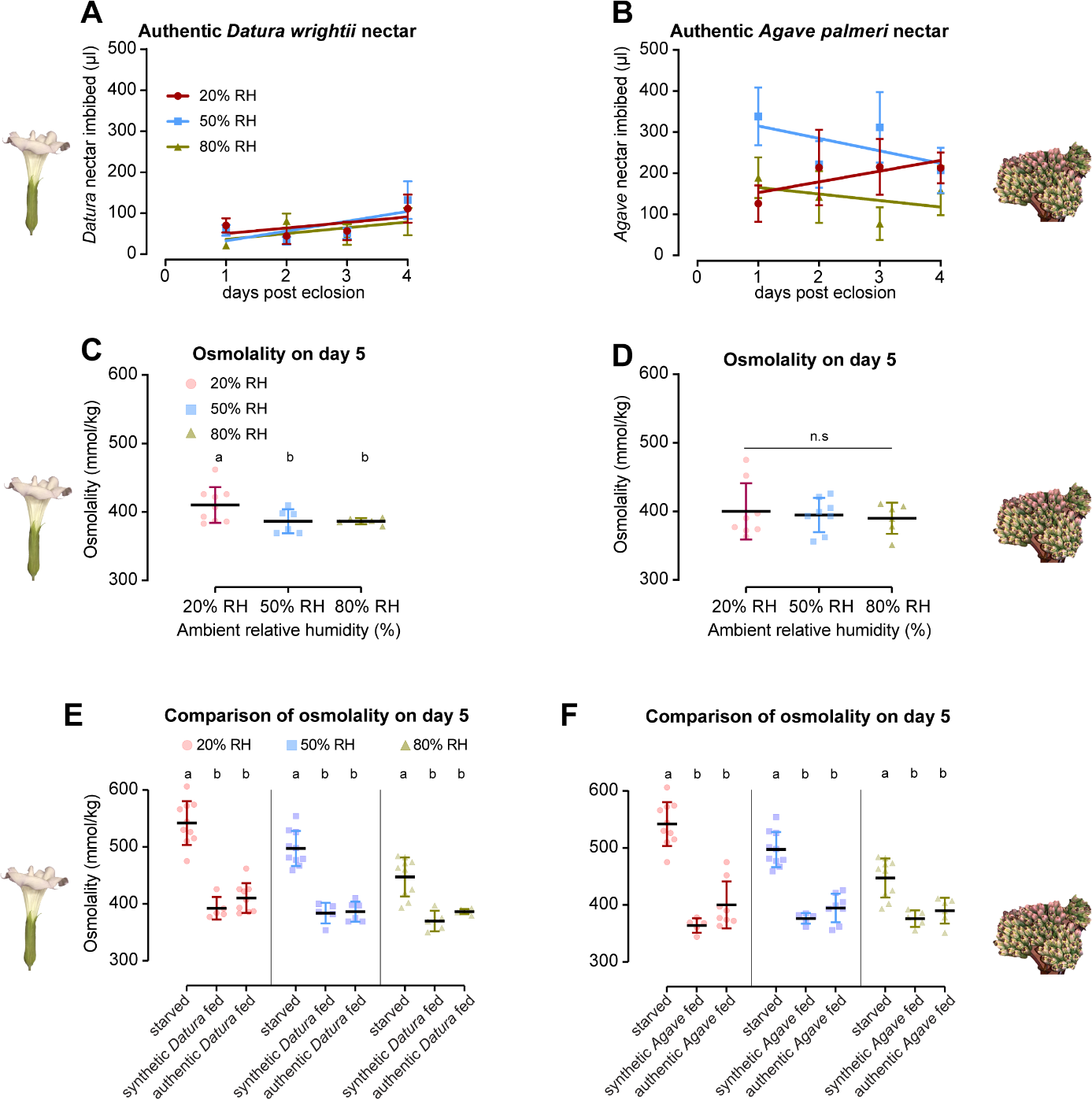
Osmolality and feeding amounts of authentic nectar-fed moths. The amount of *Datura* (A) and *Agave* nectar (B) fed by n=5-9 moths from day 1 through 4. Points show the mean and error bars show S.E. The linear regressions for feeding amounts are compared between the different RH levels for the same diet. (E, F) The osmolality of *Datura* and *Agave* nectar fed moths on day 5 post eclosion. An ANOVA is used to compare the osmolality of moths within the different RH levels on the two nectar diets. n.s indicates no significant difference among groups. Individual color-coded data points for each ambient RH level with overlaying mean (black horizontal line) and S.D have been shown. (G, H) Comparison of hemolymph osmolality on day 5 post eclosion between the authentic nectar-fed moths, synthetic nectar-fed moths (from Fig 4) and starved moths (from Fig. 2). Different letters above the figures indicate significantly different groups identified using a Fisher’s LSD test. Common letters indicate no significant difference. Data show individual color-coded data points for each condition and ambient RH and horizontal black lines show the mean with color-coded S.D.

**Fig. 6.**
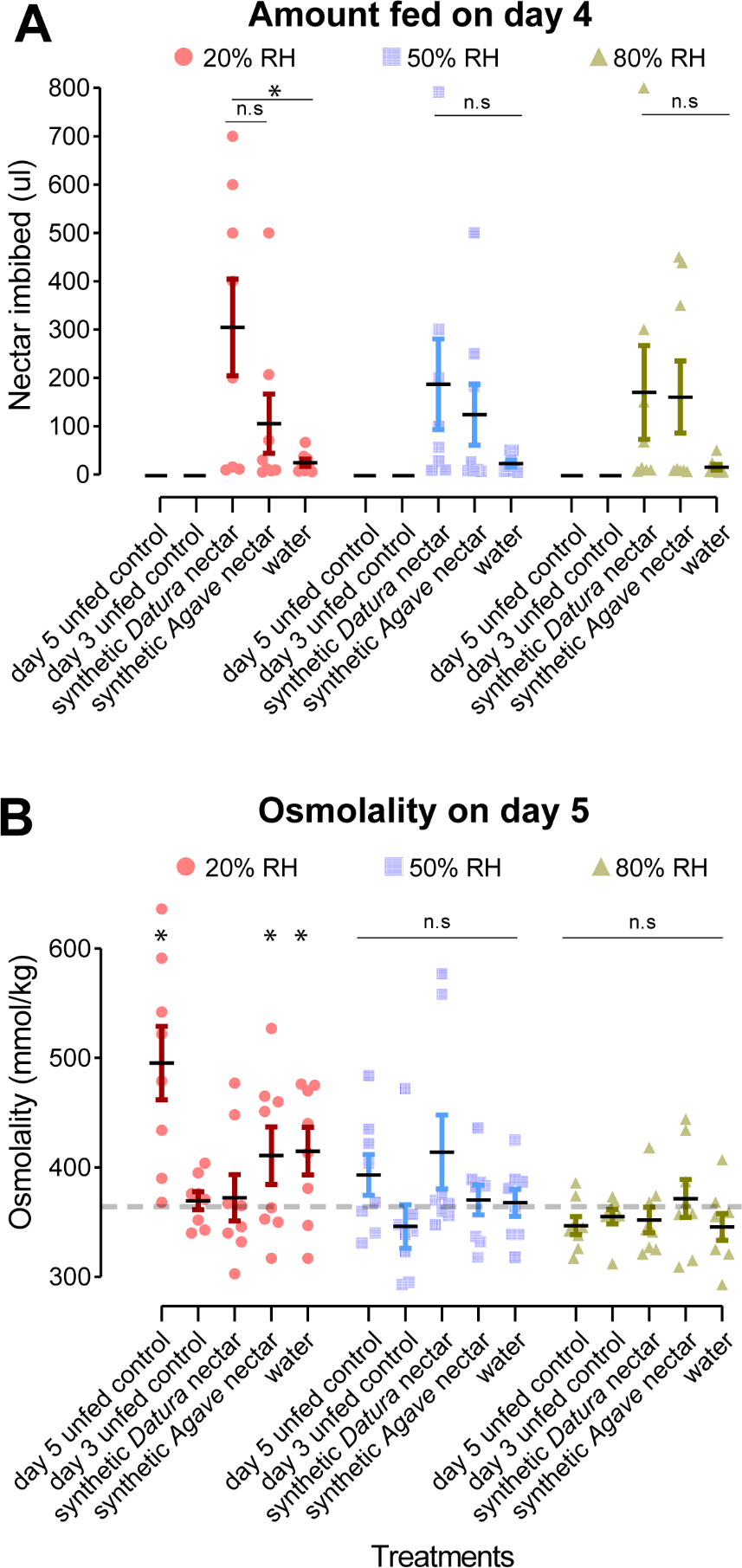
Results from the starvation-rescue experiment. Moths were fed only on day 4 to evaluate whether one nectar meal can rescue moths from osmotic stress at different ambient RH levels. As a control, the hemolymph osmolality of unfed moths was measured on days 3 and 5. Panel A shows the feeding amounts and panel B shows the corresponding hemolymph osmolality. The dashed grey horizontal line represents a baseline osmolality of 359.6 mmol/kg which is the average of day 3 unfed moths across all ambient RH levels. Data are separated and color-coded by the ambient RH level at which moths were held. Black horizontal lines represent the mean and error bars show S.E. * indicates P<0.05 and ** P<0.01, n.s. indicates no significant difference compared to the baseline osmolality.

**Table 5.** One-tailed one-sample *t*-test of moth osmolality compared against the baseline mean osmolality of 359.6 mmol/kg (average of day 3 unfed control across the different RH levels).

| Ambient RH | Individual comparisons | *t* | d.f. | *P*-value |
| --- | --- | --- | --- | --- |
| 20% RH | Starved day 5 unfed control | 4.11 | 7 | **0.0022** |
|  | Starved day 3 unfed control | 1.54 | 7 | 0.0833 |
|  | Synthetic *Datura* nectar-fed | 0.72 | 7 | 0.2453 |
|  | Synthetic *Agave* nectar-fed | 2.05 | 7 | **0.0397** |
|  | Water-fed | 2.67 | 7 | **0.0159** |
| 50% RH | Starved day 5 unfed control | 1.79 | 7 | 0.0578 |
|  | Starved day 3 unfed control | 0.67 | 7 | 0.2592 |
|  | Synthetic *Datura* nectar-fed | 1.61 | 7 | 0.0751 |
|  | Synthetic *Agave* nectar-fed | 0.78 | 7 | 0.2288 |
|  | Water-fed | 0.66 | 7 | 0.2637 |
| 80% RH | Starved day 5 unfed control | 1.25 | 7 | 0.0807 |
|  | Starved day 3 unfed control | 0.67 | 7 | 0.2596 |
|  | Synthetic *Datura* nectar-fed | 0.64 | 7 | 0.2709 |
|  | Synthetic *Agave* nectar-fed | 0.68 | 7 | 0.2577 |
|  | Water-fed | 1.14 | 7 | 0.1458 |

## Discussion

There is growing concern over the ability of pollinators to cope with climate change and the resulting shifts in floral phenology (Willmer, 2012; Scaven & Rafferty, 2013; Vanbergen & Initiative, 2013; Straka & Starzomski, 2014). The rapid decline in insect populations due to both human activity and natural factors has emphasized the importance of monitoring the health of pollinators (Janzen & Hallwachs, 2019; Sánchez-Bayo & Wyckhuys, 2019; Wagner, 2020). One potential driver of insect declines is an ecological asynchrony between flower-visiting insects and their preferred floral hosts. This phenological mismatch is especially detrimental to the survival of migrating insects that rely on food sources at stopover sites or the arrival of monsoon rains and new leaf flush at their final destinations (Srygley et al., 2010; Srygley et al., 2014; Saunders et al., 2019). In this light, it is critical to establish measurements of pollinators’ internal state as a baseline to compare starved vs. satiated or healthy vs. diseased conditions.

Here, we employed an osmometer to track the blood-water homeostasis of the nocturnal hawkmoth pollinator *Manduca sexta*, at various ambient relative humidity (RH) levels under both fed and starved conditions. The impact of dehydration stress on starved *Manduca sexta* is reflected in hemolymph osmolality measurements. Within 5 days, the osmolality of moths increased by 50%, for moths in 20% or 50% RH environments and by 39% in 8 days for those in 80% RH. At such high osmotic concentrations, moths show impaired motor control. They walk more slowly than normal, and their wings are arrested at an upstroke, indicating an imbalance of electrolytes, like muscle cramps in humans (personal observation). Moths exposed to humid environments (80% RH) survived twice as long as those in dry environments (20% RH, Fig. 2 C, D) consistent with our previous findings (Contreras et al., 2013; Contreras et al., 2022). In contrast with the starved condition, nectar or water-fed moths were able to osmoregulate by keeping hemolymph concentration within the 350-400 mmol/kg range. Our findings suggest that hemolymph osmolality could serve as an indicator of whether pollinators are food-deprived or satiated. Below we compare our findings with the hemolymph osmolalities of other insects while drawing attention to the need for establishing systematic baselines for different internal states as reference points.

There is limited research on the baseline hemolymph osmolality of pollinators. A study conducted in central Saudi Arabia tracked the hemolymph osmolality of honeybee workers (*Apis mellifera*) from day 1 to 25 in the spring and summer seasons (Ali et al., 2017). In a span of 25 days, the hemolymph concentration increased from a baseline of 330-360 mmol/kg to 490-540 mmol/kg in fed individuals, suggesting a shift in the baseline. Similar findings were observed for *Apis mellifera* drones, where the osmolality of 1-day-old drones was 334 ± 41 mmol/kg, but increased to 532 ± 38 mmol/kg for 25-day-old individuals (Leonhard & Crailsheim, 1999). These studies attribute the change in osmotic concentration to the insects’ changing behavior. The worker bees typically stay within the colony for the first 15 days after which they begin foraging outside the colony (Seeley, 1995; Ali et al., 2017). Similarly, the increase in the osmolality of the drones corresponds with their exploratory and mating flights (Leonhard & Crailsheim, 1999). In our feeding experiment, we did not find evidence of a shift in osmolality across the 8 days for *M. sexta*, which is expected as moths do not change their roles with age the way social insects do (Fig. 4). Despite the shift in baseline osmotic concentration of the honeybees sampled in Saudi Arabia, the authors concluded that the bees’ osmolality in the summer season was significantly higher than in the spring season, which indicates that in addition to the changes in age, desiccation stress is also reflected in their physiology (Ali et al., 2017). The importance of baselines is furthermore apparent in one study where wild-caught insects were sampled in the field and subsequently in a controlled lab environment (Riddle, 1986). The osmotic concentration of the Red milkweed beetle (*Tetraopes tetrophthalmus*) was 549.3 ± 16.1 mmol/kg (mean ± 95% confidence interval) in the field but decreased to 398.9 ± 29 mmol/kg after 12-16 hours of rehydration in the lab on wet paper towels. For the dogbane beetle (*Chrysochus auratus*) field measurements were 494.7 ± 10.4 which decreased to 397.8 ± 50.2 after rehydration in the lab (Riddle, 1986). Therefore, osmolality measurements can provide a physiological readout for the dehydrated vs. hydrated state of insects, whether or not they serve as pollinators.

In Table 6, we summarize the findings on the hemolymph osmolality of other insects where information about their internal state was available. The insect species in this list differ in weight over 2 orders of magnitude and present a wide range in their baseline osmolality (325 to 722 mmol/kg). Evidently, there is enormous variation in the capacity of different insects to tolerate an increase in hemolymph osmotic concentration. For example, in the Namib Desert, the beetles *Onymacris plana* and *O. unguicularis* show an increase of only 16% and 25%, respectively, from baseline osmolality post several days of severe desiccation stress (Nicolson, 1980). On the other hand, the osmolality of the Protea beetle (*Trichostetha fascicularis*) in South Africa increased by 63% from a baseline of 474 to 772 mmol/kg during a six-day period of dehydration, corresponding to a 26% decrease in body weight (Fielding & Nicolson, 1980). Species from two genera of South African hide beetles (Trogidae) show much-elevated baseline hemolymph concentration compared with other insects (Le Lagadec et al., 1998). The desert-living species *Omorgus freyi* and *O. asperulatus* lost 50% of their body weight with a corresponding increase of 86% and 70% in osmolality, respectively, from baseline. In contrast with the desert beetles, their close relatives from arid and semi-arid habitats, *O. squalidus* and *O. radula* lost 30-50% of their body weight with a corresponding increase of 22% and 30% over baseline osmolality, respectively. For smaller insects (<5 mg), such as the fruit fly and the yellow fever mosquito, an increase of only 13% and 18% over baseline osmolality reflects a stressed state. As such, the ability of insects to tolerate a large difference in osmolality is not merely a reflection of their body weight, or their habitat. Hemolymph osmolality is likely influenced by many factors, among which, water loss rates, metabolic rates, diet, and behavioral adaptations are discussed below with examples.

**Table 6.** A literature review summarizing the hemolymph osmolality of adult insects in decreasing body weight for satiated and desiccated states. The desiccation conditions in the different studies are summarized using vapor pressure deficit (VPD) as a metric to account for the differences in temperature. When the data were not explicitly mentioned, they were inferred from the text and figures in the manuscript and are indicated below with an asterisk *.

|  |  |  |  |  |  |  |  |
| --- | --- | --- | --- | --- | --- | --- | --- |
| Insect | Mean fresh mass (mg) | VPD (kPa) | No. of days dehydrated | Reported mean osmolality in different physiological conditions (mmol/kg) |  | % increase in osmolality from baseline | Reference |
|  |  |  |  | Satiated | Starved and desiccated |  |  |
| *Manduca sexta* (Tobacco hawkmoth) | 2000 | 2.4 | 5 | 360 | 541 | 50 | Present study (Data shown for males in 20% RH) |
| *Megetra cancellate* (Black and red blister beetle) | 1000* | 3 | 1 | 417 | 447 | 7 | (Cohen, 1984) |
| *Trichostetha fascicularis* (Protea beetle) | 864 | 2.95 | 6 | 474 | 772 | 63 | (Fielding & Nicolson, 1980) |
| *Onymacris plana* (Darkling beetle) | 864 | 2.95 | 12 | 429 | 498 | 16 | (Nicolson, 1980) |
| *Periplaneta americana* (Cockroach) | 825 | 1.30 | 8 | 413 | 459 | 11 | (Hyatt & Marshall, 1977) |
| *Teleogryllus oceanicus*<br>(Black field cricket) | 624 | 1.30 | 3 | 391 | 572 | 46 | (Hyatt & Marshall, 1978) |
| *Omorgus freyi*<br>(Hide beetle) | 550 | 3.4 | 10 | 568.5 | 1059 | 86 | (Le Lagadec et al., 1998) |
| *Onymacris unguicularis*<br>(Fog basking beetle) | 500 | Outdoor study | 5* | 467 | 586 | 25 | (Cooper, 1982) |
| *Omorgus squalidus*<br>(Hide beetle) | 420 | 3.4 | 3.5 | 722 | 939 | 30 | (Le Lagadec et al., 1998) |
| *Omorgus asperulatus*<br>(Hide beetle) | 400 | 3.4 | 10.5 | 635 | 1082 | 70 | (Le Lagadec et al., 1998) |
| *Omorgus radula*<br>(Hide beetle) | 200 | 3.4 | 5 | 670 | 820 | 22 | (Le Lagadec et al., 1998) |
| *Eleodes hispilabris*<br>(Desert stink beetle) | Not provided | Outdoor study | 3 | 432 | 554 | 28 | (Riddle et al., 1976) |
| *Aedes aegypti*<br>(Yellow fever mosquito) | 2.5 | 0.89 | 18 hours | ~325 (blood-fed) | ~380 | 18 | (Holmes et al., 2023) |
| *Drosophila melanogaster*<br>(Fruit fly) | 0.5 | 3.1* | 8 hours | 353 | 400 | 13 | (Albers & Bradley, 2004) |

There are several ways in which insects maintain water balance (see O’Donnell (2022)). While water can be lost through processes such as excretion, respiration (Chown, 2002), and cuticular evaporation (Gibbs, 2011), it can be gained through feeding, absorption of dew and fog (Broza, 1979; Seely, 1979), and metabolic processes (Willmer & Stone, 1997). Some insects have adaptations that allow them to lose less water through cuticular evaporation than they gain through their high metabolic processes, resulting in minimal water deficit (Nicolson & Louw, 1982). Evidence for this comes from contrasting studies on small and large carpenter bees where the smaller bees experience water deficit during foraging, but the larger bees produce excess water through their high metabolic activity (Willmer, 1988). A field study in Israel reports that the osmolality of the smaller carpenter bee *Xylocopa sulcatipes* (376-422 mg) was 483 mmol/kg before leaving the nest for foraging, and 501 mmol/kg while visiting the flowers, indicating that bees experience a water deficit. The osmolality of the larger carpenter bee *Xylocopa pubescens* (536-642 mg), however, was 529 mmol/kg before leaving the nest, and 526 mmol/kg during foraging, indicating no water deficit. In fact, the larger bees were often found excreting on flowers just before takeoff (Willmer, 1988; Nicolson, 1990).

Given the large size (10 cm wingspan), weight (2000 mg), and high metabolic activity of *Manduca sexta* it was predicted that they may experience little water deficit (Heinrich & Casey, 1973). On the contrary, the water loss rate for living moths was found to be high, with over 50-60% of their body weight lost from eclosion to senescence, even when ambient RH is high (Fig. 3). This suggests that cuticular evaporation and respiration, combined, constitute a major pathway for water loss in *M. sexta*, despite anatomical adaptations to prevent water loss. Such massive loss of water during their energetic flight must be compensated by drinking enough nectar to maintain blood-water homeostasis. An average size *M. sexta* can drink 1/3 of its body weight in aqueous fluid (∼600 µl) per day (Contreras et al., 2013). However, feeding on nectar types of different sugar concentrations poses other osmoregulatory demands for pollinators.

Nectar is not merely a blend of different sugars, but can also contain amino acids, polyamines, salts, free ions, alkaloids, and phenols (Baker & Baker, 1973; Raguso, 2004; González-Teuber & Heil, 2009; Nicolson, 2022) as well as a rich microbiome (von Arx et al., 2019). *Manduca sexta* moths exhibit an inherent preference for sugar laced with amino acids and visit artificial flowers with amino acid-augmented sugar, when those nectars also satisfy their caloric needs (Broadhead & Raguso, 2021). It is well-documented that wild-caught *M. sexta* in the Santa Rita Experimental Range near Tucson, Arizona, use both *Datura* and *Agave* flowers as major nectar sources (Alarcón et al., 2008; Riffell et al., 2008; Alarcón et al., 2010). *Datura wrightii’s* nectar sugar concentration is nearly twice that of *Agave palmeri*, and both nectars differ significantly in their sugar composition (Fig. 1). In our previous lab-based study (Contreras et al., 2013), we had only used sucrose to model the nectar sugar concentration of the sucrose-rich *Datura wrightii* (22% w/w) and the hexose-rich *Agave palmerii* (12% w/w) nectar. In this study we prepared synthetic blends to match not only the sugar concentration but also the different sugar compositions (Fig.1). The difference in hexose:sucrose ratio of the two nectars likely imposes different osmotic stress on moths. The authentic nectar of *Datura wrightii* has an osmolality of 1421.50 ± 391.26 mmol/kg (n=8), whereas its synthetic sugar mix is 1158.20 ± 100.97 mmol/kg (n=5). Similarly, the osmolality of *Agave palmeri* nectar is 924.20 ± 7.83 but its synthetic sugar mix is 818.75 ± 5.53 mmol/kg (n=4). The higher osmotic potential of authentic nectar is likely due to its amino acid, polyamine, and free ion composition (G.T. Broadhead, W. Kandalaft, R.A. Raguso, unpublished data). In our experiment, moths consumed twice as much authentic *Agave* nectar as *Datura* nectar, but overall, they consumed less authentic nectar than the synthetic sugar solutions. This disparity in consumption likely reflects the qualitative difference between the two types of nectar (authentic vs. synthetic) and moths’ osmoregulatory needs. In addition to nectar, we also fed moths with water (0% sugar). While nectar is high in osmotic concentration, water is a null solution (0 mmol/kg), posing, instead, a hypo osmotic challenge. Moths in the 20% RH condition consumed significantly more water than those in the other two ambient humidity levels, which supports previous observations of insects drinking from water puddles on dry hot days or starved moths extending their proboscis to humid air (Janzen, 1987; Raguso et al., 2005). However, water consumption itself was not sufficient for moths to maintain longevity. This was noted by the increased mortality observed within the water-fed group of moths, which was especially prominent beyond day 5 post eclosion (Fig. S1). Despite differences in nectar composition (including osmotic pressure), and the amounts consumed, well-fed moths maintain osmolality within a healthy range (350-400 mmol/kg; Fig. 5). Our findings, along with previous research on the behavioral flexibility of wild-caught moths feeding on the two nectar types (Riffell et al., 2008), highlight the physiological capacity of this large, nocturnal pollinator to balance osmotic and caloric imperatives in a semiarid habitat.

Water balance extends throughout the lifetimes of all pollinators, but what if preferred floral resources are not available for long periods of time? A forensic analysis of proboscis-borne pollen loads of *M. sexta* performed in the SRER in southeastern Arizona suggests that up to 73% of wild-caught moths may not have fed from their preferred host *D. wrightii*, 34% did not feed from their alternative host *A. palmeri,* and 15% did not feed from any flower (Levin et al., 2016). Can single bouts of nectar or water meals rescue pollinators if there are mismatches in plant-pollinator phenology? (Hegland et al., 2009; Forrest, 2015; Gerard et al., 2020). Studies predict that generalist pollinators may be better equipped than specialists to deal with mismatches in phenology (Forrest & Thomson, 2011; Kudo & Ida, 2013; Vazquez et al., 2023). In our study, we investigated the impact of an experimental 4-day mismatch between moth emergence and food availability, and whether a single nectar meal could restore osmotic balance in moths. The osmolality curves of starved moths (Fig. 2A, B) indicated an inflection point on day 4, as evidenced by the rapid increase in osmotic concentration in moths exposed to 20% and 50% RH compared to 80% RH. This increase in osmolality is particularly steep for male moths in dry environments. Therefore, day 4 is a critical point when moths must obtain a nectar meal to prevent osmotic stress. Previous work has shown that *M. sexta* flight muscle ultra-structure reaches peak maturity on day 3, (Wone et al., 2018) which coincides with their need to fly and find nectar on day 4. Our results demonstrate that a single nectar meal, or even water alone, can restore osmotic balance in moths, although we did not test its effects on longevity or fecundity. However, one or two more days of starvation could result in the death of moths if the ambient humidity is below 50% and no other food source is available. This estimate of a 4-day critical period is conservative for a habitat like the Sonoran Desert where normal daytime temperature can reach 40°C accompanied by very low RH (10% RH; VPD= 6.63 kPa)

The different ways in which insects deal with dehydration has been recently reviewed in depth (Benoit et al., 2022). An abrupt decrease in hemolymph volume or a change in electrolyte balance can cause osmotic stress and result in transcript level changes in many genes. Several hormones have been implicated in maintaining homeostasis in insects (see book chapter by Schooley et al. (2012). For example, in *Drosophila melanogaster* flies, the neuroendocrine peptide Corazonin has been shown to inhibit the release of CAPA, a diuretic hormone originally identified in *M. sexta*, which acts on the principal cells of the Malpighian tubules of flies and affects water balance (Davies et al., 2013; Terhzaz et al., 2015; Zandawala et al., 2021). In the mosquito *Aedes aegypti*, dehydration led to significant changes in the expression levels of a conserved water channel gene aquaporin 2 in the insect midgut (Holmes et al., 2023). Such findings call for proteomic or metabolomic studies of insects that are well-adapted to desiccating conditions and compare with the less adapted insects to identify novel pathways involved in desiccation resistance.

One limitation of our study is that it only focuses on virgin *M. sexta*. It is important to note that feeding rates and hemolymph osmolality may change post-mating, which represents a different internal state, especially for females. Nectar feeding significantly increases longevity and fecundity in hawkmoth females (von Arx et al., 2013), and a significant amount of body water is allocated to the developing oocytes (Nijhout & Riddiford, 1979) with mature eggs comprised of 70% water by weight (Kawooya & Law, 1988). In a study by Riddle (1986), gravid beetles showed higher hemolymph osmolality than non-gravid beetles. Our experimental design restricted moths to a specific diet, but a previous experiment with similar treatments allowed freely foraging moths to choose between all three diets simultaneously (Contreras et al., 2013). Moth activity in the different humidity levels was not documented, but wing wear and tear indicated that moths likely remained quiescent in the low (20%) and high (80%) RH environments, whereas at intermediate RH (50%), they appeared to show substantial wing damage (not quantified). In a desiccation experiment comparing the activity of *D. melanogaster* and the cactophilic *D. arizonae*, the common fruit fly showed increased activity within the first 5 hours of desiccation and no activity after 10 hours of desiccation (Gibbs, 2002). In contrast, the cactophilic *Drosophila* showed no activity during the initial hours of the experiment but became active between 10-20 hours from the start of the experiment.

Such observations highlight the different behavioral strategies used by xeric vs mesic habitat-dwelling insects. Light trapping studies in Costa Rica suggest that hawkmoths show long-distance dispersal flights, leaving the lower-elevation dry forest habitats for rainforest and cloud forest habitats (Janzen, 1987). Janzen (1986, 1987) considered various factors that could trigger such dispersal events, including seasonal shifts in host plant quality and predator pressure. Initially, we considered that plunging ambient humidity might compel moths to leave a drying habitat. However, our findings suggest that *M. sexta* moths respond to low ambient RH by becoming quiescent, as if to conserve energy until conditions improve, rather than showing increased activity that might indicate a dispersal drive (also see(Contreras et al., 2013; Contreras et al., 2022)). Such conditions inevitably led to early death. The question of which additional ecological factors, including floral nectar availability or increased perception of predation risk (Janzen, 1987), trigger hawkmoth dispersal out of dry habitats will require both field and lab assays to address. Interestingly, starved female *M. sexta* that mated with fed males produce more eggs than starved females mated with starved males, although the males do not transfer nectar-derived nutrients to the females. Instead, it is suggested that fed males provide a (neuroendocrine?) signal to the females that nectar sources are close by stimulating oogenesis in the females (Levin et al., 2016; Sendova-Franks & Foster, 2016).

Our results demonstrate the physiological resilience of a nocturnal pollinator inhabiting an environment with high desiccation stress. To evaluate the impact of anthropogenic stressors on pollinator health, standardized biomarkers have been established (reviewed in(Lopez-Uribe et al., 2020)). However, some of these techniques require molecular approaches, costly reagents, and complex post-processing for data interpretation. Hemolymph osmolality has been employed to assess the physiological status of insects, yet studies that systematically establish baseline osmolality levels for insect pollinators under different conditions are scarce. Similar data on baseline hemolymph osmotic concentrations for insects affected by parasites (Jones et al., 2022), fungal pathogens (Evison & Jensen, 2018), or exposed to pesticides and insecticides (Goulson et al., 2015), in comparison to healthy control populations, would provide valuable resources to track pollinator health . Therefore, controlled lab experiments could serve as a reference to rapidly monitor the health of wild insect pollinators in their natural habitats. Such resources can inform policymakers and conservationists to implement quick and appropriate restoration efforts.

Climate change is causing a mismatch in the timing of pollinators and flower phenology. The effects of this mismatch are likely to be particularly acute in desert ecosystems where precipitation events are more likely to be disrupted (NOAA Drought Task Force, 2021). This study shows that *Manduca sexta* moths can tolerate a mismatch in available nectar for about three days (half their lifespan), after which they exhibit extreme osmotic stress. This osmotic stress is strongly dependent on the changes in relative humidity that co-occur with precipitation events. Fortunately, for these important desert pollinators, a single nectar meal can mitigate the effects of osmotic stress of starved moths imposed by the mismatch in nectar resources.

## Acknowledgments

We would like to thank Nathan Oakes for setting up the closed-loop humidity monitoring system in the incubator. We thank Lynn Marie Johnson from Cornell’s statistical consulting unit for helping with the ANOVA models. This work was funded by National Science Foundation Grants IOS-0923765 (to J.G. and R.A.R.) and IOS-0923180 (to G.D.). Parts of this research were funded by the Cornell Agriculture and life science alumni grant and Cornell Sigma Xi awarded to A.D.

